# Horizontal transmission of the entomopathogenic fungal isolate INRS-242 of *Beauveria bassiana* in emerald ash borer, *Agrilus planipennis* Fairmaire

**DOI:** 10.1101/532838

**Authors:** Narin Srei, Robert Lavallée, Claude Guertin

## Abstract

Emerald ash borer (EAB), *Agrilus planipennis* Fairmaire, is an invasive and destructive beetle that causes extensive damage to ash trees in North America. The entomopathogenic fungus *Beauveria bassiana* is considered as an effective biological control agent for EAB adult populations. Using an autodissemination device with a fungal isolate of *B. bassiana*, our research aims to investigate the possibility of horizontal transmission of the fungal disease from infected to uninfected EAB adults during mating. Results show that the efficiency of fungal transmission is significantly related to the sex of EAB carrying the fungal pathogen. EAB males are the promising vector to transmit mycosis to their partners during mating. Results strengthen the potential of the fungal autodissemination device as a powerful biological strategy to control EAB populations.

## Introduction

Emerald ash borer (EAB), *Agrilus planipennis* Fairmaire, is an extremely invasive insect that is responsible for killing hundreds of million ash trees, *Fraxinus* spp., in urban and forest ecosystems of North America. EAB was first detected in 2002 in Detroit, Michigan (USA) and in Windsor, Ontario (Canada) after its introduction from northeastern Asia (Haack et al. 2002). To date, EAB is found in 35 American states and five Canadian provinces (Emerald ash borer information network). After hatching, EAB larvae tunnel through the bark and begin feeding the phloem and the cambium (Cappaert et al. 2005). The abundance of larval galleries disrupts the flow of nutrients and water within the tree and may cause the tree death in 3-4 years (Siegert et al. 2009).

Various strategies have currently developed to suppress EAB populations. These approaches include the use of systemic insecticides (Herms et al. 2014), parasitoids (Duan et al. 2012) and fungal control agents (Lyons et al. 2012). However, management of EAB is still challenging because of the beetle’s cryptic behaviors. A fungal autodissemination device developed by Dr. Guertin and Dr. Lavallée has shown its potential in controlling EAB (Lyons et al. 2012). Results shown that most EAB adults who contacted the autodissemination chamber died after five days. This study builds upon previous research and aims to test the hypothesis that infected adults released from the autodissemination device can horizontally transmit the fungal pathogen to uninfected partners during mating.

## Materials and Methods

### Emerald ash borer Collection

During summer of 2017, six 12-funnel green Lindgren traps (Synergy Semiochemicals, Burnaby, BC, Canada) were suspended in the upper third of the canopy of ash trees located in Laval, QC, Canada (Long. 45.541309; Lat. −73.718103). EAB adults captured in the collecting jars were harvested daily. In the laboratory, male and female EAB were separated using morphological characters (Rodriguez-Saon et al. 2007). Insects were then reared in cages and fed with fresh ash leaves until their use in experiments.

### Fungal production

The isolate INRS-242 of *Beauveria bassiana* (also known as INRS-CFL), which is an effective entomopathogen of *A. planipennis* adults (Lyons et al. 2012), was used in this study. This isolate is an endogenous fungus recovered by Lavallée and Guertin from the pine shoot beetle, *Tomicus piniperda* L., near Cookeshire, QC, Canada. The fungal isolate is stored in 70% of glycerol at −80°C in the Fungal bank of the INRS-Armand-Frappier Santé and Biotechnologie, Laval, QC, Canada.

Autodissemination devices containing fungal coated-pouch were used in this experiment to inoculate EAB males and females. The production of the pouches was achieved as described by Srei (2017). Briefly, the INRS-242 isolate was first grown in Yeast Extract Peptone Dextrose broth (YPD, Alpha Bioscience Inc., Baltimore, MD, USA), in a rotary shaker (150 rpm) for four days at 25°C. The pouches (14.5 x 11 cm; fiberglass mosquito net) containing sterile pearled barley were then inoculated with the fungal conidial suspension before incubation for ten days in the growth chamber under controlled conditions (25°C, 70% of humidity and in the darkness).

### Experimental design

A randomized block design with three replicates (blocks) was used to assess the transmission of the fungal disease from infected to uninfected EAB adults during mating behavior. Three treatments were randomly assigned to each block, with 20 EAB pairs per treatment. The first treatment represents the control pair, in which one uninfected male and one uninfected female were introduced in an arena (Petri Dishes 100 x 150 mm, Fisher Scientific, Ontario, Canada). For the second treatment, only the EAB male was inoculated with the fungus (IM+UF), whereas in the third treatment, just the EAB female was contaminated with the fungus (IF+UM). Three sterilized pouches without fungal isolate were used for the control treatments. For each infection treatment of each block, a fungal-coated pouch was employed once to inoculate EAB adults (Srei et al. 2017). Briefly, after walked on the fungal-coated pouch for 1 min, each EAB male or female was transferred into the arena using sterilized tweezers. For all treatments, each EAB female was glued to the Petri dish surface using Evo-Stik (Evode Industry Ltd., Newtown, Ireland). This approach avoids the possibility of indirect horizontal transfer resulting from insect movement inside the arena. After being placed in the Petri dishes, the insects were observed for 18 hours. Two parameters were measured during the assay: presence of mating and fungal transmission. The fungal transmission was recorded as occurring when conidia were present on the ventral side of uninfected males, or on the dorsal part of uninfected females. In addition to these parameters, the subsequent mortality of EAB was also reported. After 18 hours, EAB males and females from each arena were individually transferred to a new Petri dish in which a fresh ash leaf was provided as food substrate. All Petri dishes were then incubated in the growth chamber at 25°C with 50% of relative humidity and 16:8 h photoperiod (light:dark). Mycosis was confirmed by the appearance of white muscardine on EAB cadavers. Both mortality of EAB adults and the presence of the muscardine were recorded daily for 14 days.

### Statistical analysis

All statistical analyses were performed using the software R, version 3.4.3. The proportion of mated pairs and horizontal transmission efficacy of INRS-242 isolate of *B. bassiana* among EAB adults were assessed using an analysis of variance (ANOVA) to show significant differences between treatments. Means associated with different treatments were then coated using Duncan’s multiple comparisons. Student’s *t*-test was used to compare the fungal infection percentage of mated pairs and unmated pairs. For all statistical tests, the level of rejection was set at α = 0.05.

## Results

A significant difference in the percentage of EAB adults that mated was observed between treatments (ANOVA, df = 2; F value = 6.75; *p* = 0.0291) (Table 1). More precisely, no difference was recorded between the control group and IM+UF treatment. However, the percentage of insects that mated was significantly lower in IF+UM treatment. In both infection treatments (IM+UF and IF+UM), mycosis transmission was of 100% for mated beetles. On the other hand, no mycosis was observed in mated EAB pairs of control groups. As expected, the percentage of infected EAB was significantly higher in mated pairs than unmated pairs, with 66.7% and 4.4% respectively (*t*-test = 3.6766; df = 16; *p* = 0.002). All infected EAB adults died within six days following the fungal contamination, and white muscardine was recorded on all contaminated individuals.

**Table 1.**
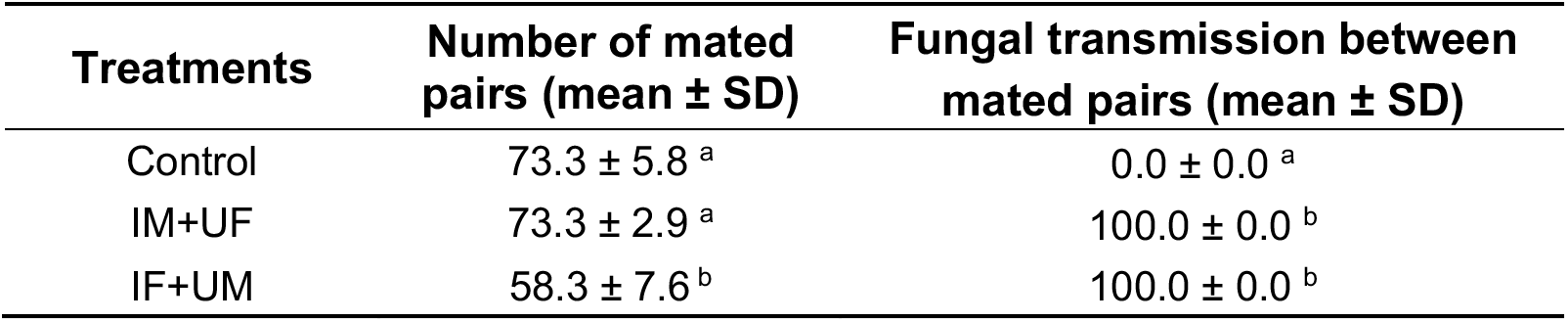
Percentage of mated pairs per treatment and the proportion of horizontal transmission of *B. bassiana* isolate INRS-242 recorded in adults of *A. planipennis* mated pairs. Control: male and female EAB were not exposed to the fungal isolate; IM+UF: fungal transmission between infected male and uninfected female EAB; and IF+UM: fungal transmission between infected female and uninfected male EAB. For each column, distinct letters indicate a significant difference (Duncan, n = 60, *p* < 0.05).

| Treatments | Number of mated pairs (mean ± SD) | Fungal transmission between mated pairs (mean ± SD) |
| --- | --- | --- |
| Control | 73.3 ± 5.8 <sup>a</sup> | 0.0 ± 0.0 <sup>a</sup> |
| IM+UF | 73.3 ± 2.9 <sup>a</sup> | 100.0 ± 0.0 <sup>b</sup> |
| IF+UM | 58.3 ± 7.6 <sup>b</sup> | 100.0 ± 0.0 <sup>b</sup> |

## Discussion

Horizontal transmission by direct contact between fungal-infected and uninfected insects is a mechanism of natural dispersion of conidia that regulates insect population size (Steinkraus 2006). This fungal transmission could occur from the time of initial contact during mating (Kreutz et al. 2004), as has previously observed in insect species including the eastern larch beetle, *Dendroctonus simplex* LeConte (Srei 2017), the European spruce bark beetle, *Ips typographus* L. (Kreutz et al. 2004), the Mexican fruit fly, *Anastrepha ludens* Loew (Toledo et al. 2007), and the stem borer, *Busseola fusca* Fuller (Maniania et al. 2011). Under laboratory conditions, our results have demonstrated for the first time that an infected EAB adult can transmit *B. bassiana* isolate INRS-242 to an uninfected mate by contact during mating. Furthermore, our results shown that no fungal transmission was recorded on insects having no contact, which provides evidence that the confinement did not mediate mycosis transfer. For all EAB mated pairs, the average duration of mating was about 42 min (data not shown). This observation corroborates to that of Pureswaran and Poland (2009). Following the mortality of EAB adults and the appearance of muscardine, it is possible to confirm that the fungal dose transmitted from infected to uninfected EAB adults during mating is strong enough to cause the death of insects.

The efficiency of fungal transmission depends on the sex of the insect that carries the pathogen. Transmission of entomopathogenic fungus *Metarhizium anisopliae* (Metschn.) Sorokin within the populations of the Mediterranean fruit fly, *Ceratitis capitata* Wiedemann, was higher when males were carriers of the pathogen (Quesada-Moraga et al. 2008). In *D. simplex*, horizontal transmission of *B. bassiana* isolate INRS-242 was greatest when females were inoculated with the fungus (Srei 2017). Our results show that male-to-female transmission of the fungal isolate within EAB adults was higher than from female-to-male. One distinct characteristic of EAB males is their pubescence. This may facilitate the adhesion of conidia and could explain the difference observed in fungal transmission between infected males and infected females. Moreover, a preliminary experiment suggests that some EAB males can mate with at least two different females (data not shown). This observation suggests that increasing the proportion of contaminated males can be a valuable strategy for increasing the mortality rate of the adult EAB population. Adding another mortality factor to other biotic and abiotic factors could contribute to slow or to keep the EAB population at a lower level and may eventually reduce oviposition on ash trees.

## Acknowledgments

We are grateful to Cynthia O’Hearn for technical support. This research was supported by 2016 GDG Environment/NRCan/INRS funding agreement to C.G. and R.L.

